# MICOMWeb: a website for microbial community metabolic modeling of the human gut

**DOI:** 10.1101/2025.03.14.643140

**Authors:** Cristóbal Fresno, Juan José Oropeza-Valdez, Perla Itzel Alvarado-Luis, Patricia Peña-González, Armando R Tovar, Nimbe Torres, Christian Diener, Sean Gibbons, Osbaldo Resendis-Antonio

## Abstract

**Summary:** MICOMWeb is a user-friendly website for easy microbial community metabolic modeling of the human gut. This website tackles three constraints when generating *in silico* metagenome-scale metabolic models: (i) the prior Python user knowledge for metabolic modeling using flux balance analysis with MICOM Python package, (ii) predefined and user-defined diets to generate *ad hoc* metabolic models, and (iii) the high- throughput computational infrastructure required to obtain the simulated growth and metabolic exchange fluxes, using real abundance from metagenomic shotgun or 16S amplicon sequencing; we present MICOMWeb’s features to easily run *in silico* experiments as a functional hypothesis generator for experimental validation on previously published data.

**Availability and Implementation:** MICOMWeb is freely accessible for academic purposes at https://micomweb.c3.unam.mx.

## 1 Introduction

Over the last decades, next-generation sequencing platforms have made it possible to move from a single to a multi-organism analysis. A single bacterial study has moved into microbial communities where the challenge is to study how population interactions (antagonism, neutralism, mutualism, etc.) propitiate their respective global ecological equilibrium (Bosi *et al*. 2017). In recent years, some computational schemes have been suggested to disentangle and discern the metabolic complexity in microbial communities, most of them assuming flux balance analysis as the fundamental mathematical core (Orth *et al*. 2010).

Currently, different challenges exist for microbial community metabolic modeling from biological and computational perspectives (Bosi *et al*. 2017). Large-scale community modeling with the associated numerical stability has been addressed by diverse approaches such as MICOM (Diener et al. 2020) and PyCoMo (Predl *et al*. 2024). On the other hand, modeling itself is a computationally intensive task, which combined with the first challenge, leads us to an additional problem: technology adoption for community metabolic modeling (Meyerovich & Rabkin 2013). In this domain-specific niche, the ruling programming language is Python (Python Software Foundation 2022); hence, the research user must know biology and learn/adopt Python if unfamiliar.

To reduce these gaps and facilitate the use of these computational models, here we present MICOMWeb, a friendly-user academic website (https://micomweb.c3.unam.mx) for easy microbial community metabolic modeling of the human gut using MICOM, a previous algorithm for modeling metabolic activity at the genome-scale level (Diener et al. 2020). The user can input the genome-scale microbiota composition of their experiment and obtain the microbiota growth rates, metabolite exchanges, and annotation out of the box without the need for a programming language learning curve and computational infrastructure requirements. For exemplification, MICOMWeb performance is presented using real data previously published by Weingarden *et al*. 2015.

## 2 Description

MICOMWeb is an academic website that assists *in silico* mechanistic hypotheses generation of microbial metabolic activity based on how diet and microbiome composition influence community functions. Its frontend (Fig. 1A) is based on the Django-Python 3 framework (Django Software Foundation 2022, Python Software Foundation 2022) and Bootstrap 5 (Bootstrap 2023) over the Secure Sockets Layer (Weaver 2006). Users can register using MICOMWeb’s sign-in or OAuth 2.0 protocol (IETF 2012) with Gmail, GitHub, and ORCID. A companion MySQL database (MySQL 2022) stores credentials and Django models.

**Fig. 1.**
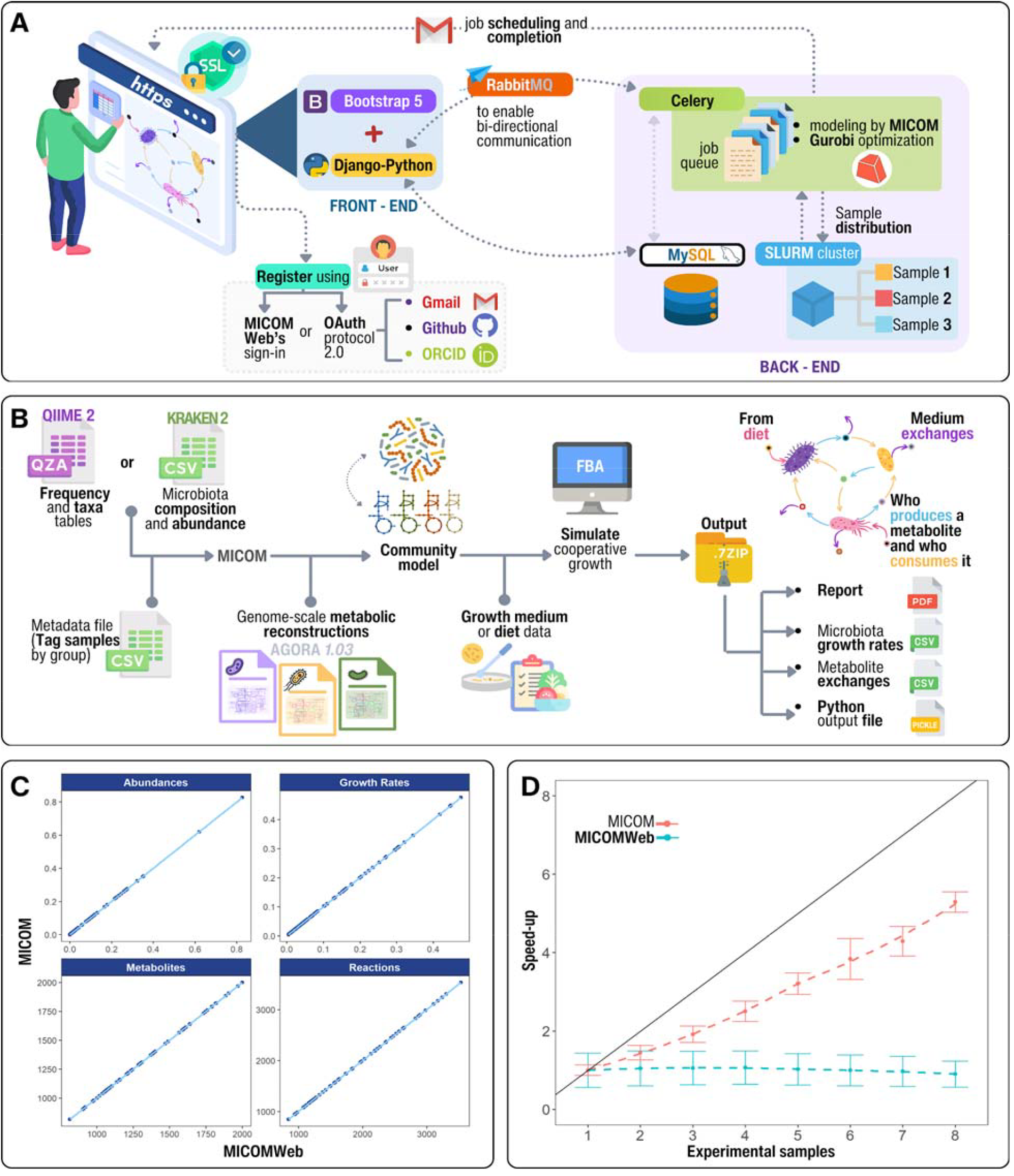
MICOMWeb overview. (A) high-level front to backend infrastructure. (B) *micr*obial *com*munity modeling with the MICOM Python package (Diener *et al*. 2020). (C) MICOMWeb vs. MICOM reproducibility comparison on abundances, growth rates, number of metabolites, and reactions. (D) The speed-up curve for the example dataset (Weingarden *et al*. 2015) using MICOM and MICOMWeb.

The bi-directional communication between the front and backend is handled by RabbitMQ (RabbitMQ 2023, Fig. 1A). The backend workload is orchestrated using the Celery job queue (Celery Project 2023). The microbial community metabolic modeling of the human gut is handled by the MICOM Python package (Diener *et al*. 2020) using Gurobi (https://www.gurobi.com/) to solve the quadratic equation system. However, to tackle the high-throughput computational requirements, MICOMWeb distributes the experimental data samples into an SLURM cluster (Simple Linux Utility for Resources Management, Jette & Wickberg 2023). The user gets email notifications on job scheduling and completion.

MICOMWeb can handle for inputs QIIME 2 (Bolyen *et al*. 2019) artifacts such as frequency and taxa table (.qza files), as well as a comma separated value files (.csv) with the microbiota composition and abundance (Fig. 1B). In either case, an additional metadata (.csv) file is required to label the samples with an experimental group tag for quality control.

The inputs are loaded into MICOM and summarized according to the taxonomic rank level (‘*genus*’ by default). The genome-scale community metabolic reconstructions per sample will be created using AGORA da 2017). Detailed information on the macronutrients in the diet and the relative abundance is sufficient to simulate cooperative growth (Diener *et al*., 2020). The cut-off value by default is at least 0.01% of the sample abundance.

Once the metabolic reconstruction models are created, the user selects a pre- or user-defined growth medium to simulate the flux analysis balance yielding the microbiota growth rates, metabolite exchanges, annotation, or even the complete Python output file (.pickle) for further analysis. In addition, MICOMWeb also offers a complete run report. Output files are stored and delivered as compressed 7-Zip files (Pavlov 2022) upon user request on the website (Supplementary data).

## 3 Results

MICOMWeb was tested using data previously published by Weingarden *et al*. (2015). Briefly, 16S rRNA gene high-throughput sequencing was performed on fecal samples from four healthy donors and four patients before fecal microbiota transplantation (FMT) to treat refractory *Clostridium difficile* infection.

The example dataset is composed of three files that contain the eight samples: i) the QIIME 2 artifacts for the frequency table (*table*.*qza*), ii) the taxa composition (*taxa*.*qza*), and iii) the metadata file (*metadata*.*csv*) to tag healthy-donor and pre-FMT samples.

The microbial community metabolic modeling was run using a cluster with 20 nodes with a total of 4,224 GB of distributed memory and 1,208 threads running Centos version 7 with InfiniBand connectivity and Lustre file system (CentOS Project 2024, InfiniBand Trade Associatio. 2017) under default parameters.

First, we validate MICOMWeb vs. MICOM reproducibility by comparing the output results obtained by each strategy in terms of abundances, growth rates, number of metabolites, and reactions. (Fig. 1C). The scatter plot panel results suggested that all the points fall into the identity line, proving that the correlation between MICOMWeb and MICOM is almost perfect, a signal of reproducibility. The companion experiment report run is available as Supplementary material.

Second, to validate MICOMWeb’s scalability, we compared ten unconstrained MICOMWeb runs with ten MICOM runs using one and eight threads (one thread per sample). The original MICOM’s implementation had an execution time of 01:38:37 ± 00:01:22 and 01:48:13 ± 00:20:55 from sequential to parallel implementation, with a total RAM consumption of 3.65 ± 0.05 and 15.35 ± 0.01 GB, respectively. In contrast, MICOMWeb distributed each sample independently per single thread and had a total computational resource consumption of 1.41 ± 0.27 GB of memory per sample and 15:10 ± 4:22 for total execution time. The reduction in computational resources between the two implementations was 10.92 folds for memory and 7.13 folds for total execution time.

Finally, we simulate both MICOM and MICOMWeb experiment size scalability constrained to full SLURM cluster availability (Fig. 1D). We simulated ten consecutive runs using all possible combinations of the FMT dataset taken simultaneously by one to eight samples, with the same number of threads as samples were taken. The speed- up-like curve results suggest that MICOM’s implementation departs from the identity behavior *y* = *x* as expected. In contrast, MICOMWeb has a constant run-time independent of the number of samples provided. In practical terms, this behavior is upper bounded by the sample with the greatest microbiota diversity, i.e., the sample with the biggest metabolic reconstruction model size.

## 4 Conclusion

MICOMWeb is an academic user-friendly website that assists in silico metabolic mechanistic hypotheses generation based on how diet and microbiome composition influence community functions. In minutes, the research community can register into the platform and be ready to upload their experimental high-throughput microbiota composition, build the metabolic reconstruction, and simulate the flux analysis balance to easily obtain the microbiota growth rates, metabolite exchanges, and annotation. Traditionally, obtaining these results would have required the researcher to have previous knowledge of Python or be forced to adopt it to use the MICOM base package. In addition, we have to consider the required computational infrastructure to run the analysis, administration, and additional related costs.

MICOMWeb was tested using 16S rRNA high-throughput sequencing data from Weingarden et al. 2015, which also showed reproducibility compared to its original Python implementation. Moreover, the web-based implementation counterpart uses a SLURM cluster to distribute the experimental samples’ flux balance analysis workload without the researcher’s intervention, except when receiving the completion email to download the results.

Finally, MICOMWeb, as a companion academic website for microbial community metabolic modeling of the human gut, opens new research in testable experimental metabolic mechanistic hypothesis generation horizons that would not have been feasible to perform *in vivo*.

## Acknowledgments

The authors would like to acknowledge), in particular to José Luis Gordillo-Ruiz from the High-Performance Computing Unit (HPCU) for proving the high availability of cluster infrastructure required for this project. PIAL is a student of the MSc program at the Maestría en Ciencias Bioquímicas, Universidad Nacional Autónoma de México, and received a CONAHCyT fellowship (CVU 1273128). CF had a postdoctoral fellowship for this project funded by the National Council of Humanities, Sciences, and Technologies (CONAHCyT) [Grant Ciencia de Frontera 2019, FORDECYT-PRONACES/425859/2020]. JJ-OV thanks the financial support from the UNAM Postdoctoral Program DGAPA.

## Data availability

The microbiota example data used in this article is freely available under BioProject PRJEB19996 (https://www.ncbi.nlm.nih.gov/bioproject/PRJEB19996) and QIIME2 artifacts at MICOMWeb’s website (https://www.micomweb.c3.unam.mx). MICOM open-source package is available using the Python package management system (pip) and at the developer website (https://github.com/micom-dev/micom). Quadratic problems were solved using Gurobi (https://www.gurobi.com/) academic site license for research purposes.

## Supplementary data

Supplementary data that contains MICOMWeb’s outputs, growths, KOs, and reports can be found on Mendeley data (https://data.mendeley.com/datasets/zybpsy2djp/1, doi: 10.17632/zybpsy2djp.1).

